# Belgian *Culex pipiens pipiens* are competent vectors for West Nile virus but not Usutu virus

**DOI:** 10.1101/2023.05.17.541091

**Authors:** Alina Soto, Lander De Coninck, Ann-Sophie Devlies, Van De Wiele Celine, Ana Lucia Rosales Rosas, Lanjiao Wang, Jelle Matthijnssens, Leen Delang

**Author notes:** Corresponding author: Leen Delang.

## Abstract

**Background:** West Nile virus (WNV) and Usutu virus (USUV) are emerging arboviruses in Europe transmitted by *Culex* mosquitoes. In Belgium, it is currently unknown which *Culex* species are competent vectors for WNV or USUV and if these mosquitoes carry *Wolbachia*, an endosymbiotic bacterium that can block arbovirus transmission. The aims of our study were to measure the vector competence of Belgian *Culex* mosquitoes to WNV and USUV and determine if a naturally acquired *Wolbachia* infection can influence virus transmission.

**Methodology/Principal Findings:** We captured 876 non-engorged female *Culex* mosquitoes from urban and peri-urban sites in Leuven, Belgium. We provided females with an infectious bloodmeal containing WNV lineage 2, USUV European (EU) lineage 3, or USUV African (AF) lineage 3. Blood-fed females (n=154) were incubated for 14 days at 25°C after which the body, head, and saliva were collected to measure infection (IR), dissemination (DR), and transmission (TR) rates, respectively. Mosquito species were identified by qRT-PCR or Sanger sequencing, the presence of infectious virus in mosquitoes was confirmed by plaque assays, and viral genome copies were quantified by qRT-PCR. We found that *Culex pipiens pipiens* were able to transmit WNV (11% IR, 40% DR, 100% TR) but not USUV (EU lineage: 13% IR, 0% DR; AF lineage: 16% IR, 17% DR, 0% TR). In contrast, *Culex modestus* was able to transmit USUV (AF lineage: 60% IR, 67% DR, 50% TR), but not WNV (0% IR). We found that the presence or absence of *Wolbachia* was species-dependent and did not associate with virus transmission.

**Conclusions/Significance:** This is the first report that Belgian *Culex* mosquitoes can transmit both WNV and USUV, forewarning the risk of human transmission. More research is needed to understand the potential influence of *Wolbachia* on arbovirus transmission in *Culex modestus* mosquitoes.

**Author Summary:** West Nile virus and Usutu virus can cause seasonal epidemics in humans. They are maintained in a transmission cycle between wild birds and *Culex* mosquitoes, and humans that are bitten by infected mosquitoes can develop life-threatening neurological disease. Certain *Culex* species carry the symbiotic bacterium *Wolbachia* which can block virus transmission in mosquitoes. In Belgium, it is currently unknown which *Culex* species can transmit West Nile virus and/or Usutu virus, or if they carry *Wolbachia* bacteria. In our study, we captured wild mosquitoes from Belgium and infected them with West Nile virus or Usutu virus. We found that a common European species (*Culex pipiens pipiens*, the Northern House mosquito) could transmit West Nile virus, whereas a lesser known species (*Culex modestus*) could transmit Usutu virus. *Wolbachia* bacteria could be found in almost all *Culex pipiens pipiens*, but not in *Culex modestus*, suggesting that *Wolbachia* prevalence is species-specific. More research is needed to understand if *Wolbachia* can block virus transmission in *Culex modestus*. This is the first report on the ability of *Culex* mosquitoes to transmit West Nile virus and Usutu virus in Belgium, forewarning the risk of transmission to humans.

## Introduction

West Nile virus (WNV) and Usutu virus (USUV) are emerging arboviruses in Europe. They are both flaviviruses (Family: *Flaviviridae*) and members of the Japanese Encephalitis serocomplex, sharing considerable similarities in their transmission and clinical relevance. The lifecycle of WNV and USUV is enzootic: they amplify in resident and migratory birds and are transmitted to new hosts via intermediary mosquito vectors. Mammals, including humans, can become incidental hosts of WNV or USUV when bitten by infected mosquitoes. From 2010 to 2022, there were over 5,800 reported human cases with 378 deaths caused by WNV in Europe (1,2). In contrast, there have been few human cases of USUV detected in Europe – only 17 reports of neuroinvasive disease so far – as symptomatic infections are rarely detected (3). Cross-reactive WNV nucleic acid tests from human blood and organ donor screenings have led to the incidental identification of passive USUV cases, which suggests that the true incidence of USUV is underestimated (3). In Belgium, no human cases of WNV or USUV have been reported, but the country is considered at-risk. Neighboring countries have experienced recent human cases of WNV, with evidence of WNV RNA detection in native birds and mosquitoes in the Netherlands (4,5) and Germany (6–8), while USUV is reported endemic to resident birds and bats in Belgium since 2016 (9).

The most important vectors for WNV and USUV are members of the genus *Culex* (Family: *Culicidae*). The vectors established in Europe are *Culex pipiens* (Linnaeus 1758), *Culex modestus* (Ficalbi 1889), *Culex torrentium* (Martini 1925), and *Culex perexiguus* (Theobald 1903) (10–17), of which all but the latter are present in Belgium (18,19). *Culex pipiens* (*sensu lato*, s.l.) can be divided into two morphologically identical but behaviorally distinct biotypes: *Culex pipiens pipiens* (Linnaeus 1758) and *Culex pipiens molestus* (Forskål 1775). *Culex pipiens pipiens* is an established European vector for WNV, based on evidence from vector competence studies using field-caught mosquitoes (11,20,21). USUV RNA has been detected in native European *Culex pipiens pipiens* (22,23), but so far vector competence studies using live mosquitoes have been restricted to laboratory colonies (24,25). Of the two biotypes, *pipiens* is the more efficient vector for both WNV (21,26) and USUV (25). In field-captured *Culex modestus* mosquitoes, WNV and USUV RNA have been detected (15,27), but currently the only evidence of WNV vector competence in live mosquitoes comes from laboratory colonies (12,28). So far, there are no measures of USUV vector competence in *Culex modestus* using field or laboratory mosquitoes. Therefore, the vector competence of *Culex pipiens pipiens* to USUV, and of *Culex modestus* to both WNV and USUV, using native vectors from natural habitats have not been investigated.

The presence of *Wolbachia pipientis* in mosquitoes should be an important consideration in vector competence studies*. Wolbachia* are intracellular gram-negative alphaproteobacteria found to naturally infect most arthropod species worldwide (29). In arboviral research, *Wolbachia pipientis* are well known for their ability to reduce the fitness and reproduction of mosquitoes and suppress arbovirus transmission, particularly in *Aedes aegypti* (30). Several strains of *Wolbachia* can directly interfere with arbovirus replication in mosquitoes [reviewed by (31)], but the evidence on *Wolbachia*-mediated inhibition of WNV is contradictory - as it remains unclear if *Wolbachia* enhances or protects against WNV transmission (32–36). More than 90% of *Culex pipiens* s.l. harbor *Wolbachia* (32,37), whereas there is limited evidence that *Culex modestus* carry this bacterium (38,39). As of yet, there are no studies evaluating the influence of *Wolbachia* on arbovirus transmission in *Culex modestus*, or on USUV transmission in any species.

The aims of our study were to determine the vector competence of Belgian *Culex* mosquitoes to WNV and USUV and investigate if the presence of *Wolbachia* confers protection against transmission. We captured female *Culex* mosquitoes from urban and peri-urban sites and identified them based on morphology, molecular identification, and DNA barcoding. Next, captured mosquitoes were fed an infectious bloodmeal containing either WNV or two different USUV strains to determine infection, dissemination and transmission rates, and determined the prevalence of *Wolbachia*. This is the first report on the ability of Belgian mosquitoes to harbor *Wolbachia* and transmit WNV and USUV.

## Methods

### Mosquitoes

*Culex* mosquitoes were collected in the field from June-September 2022 in Leuven, Flemish Brabant, Belgium (Fig. 1). Collections took place interchangeably between an urban habitat (The Botanical Garden of Leuven, N 50°52’41, E 4°41’21) and a peri-urban habitat (Arenberg Park, N 50°51’46, E 4°41’01). Adult mosquitoes were captured using BG Sentinel traps (Biogents® AG, Regensburg, Germany) baited with dry ice for constant CO_2_ release and a sachet of BG-Sweetscent (Biogents® AG, Regensburg, Germany) to imitate the scent of human skin. Traps were placed in dispersed locations at either habitat in the late afternoon to allow the capture of free-flying mosquitoes overnight. Trapped mosquitoes were collected the following morning and transported to an insectary facility for sorting. Mosquitoes were anaesthetized over dry ice and identified morphologically to the *Culex* genus level. The sex and feeding condition (unfed/gravid/blood-fed) of mosquitoes were determined based on morphological cues. Females were placed in 32.5 cm^3^ BugDorm cages (MegaView Science Co., Ltd., Taichung, Taiwan) with access to 10% sucrose *ad libitum* on cotton pledgets. The cages were kept for up to one week in an incubator set to 25°C and 70% relative humidity (RH) with a photoperiod of 16:8 light:dark hours.

**Figure 1.**
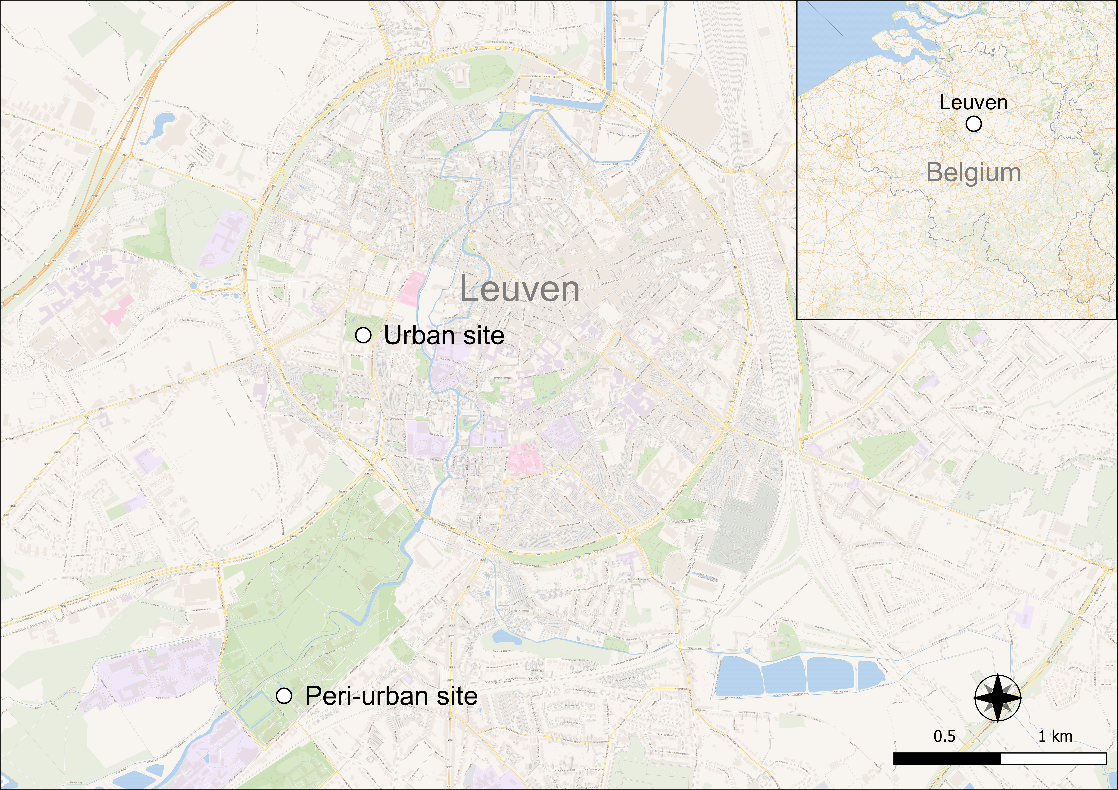
Urban and peri-urban field collection sites in Leuven, Flemish Brabant, Belgium. Map made using QGIS software.

### Cell Lines & Virus Stocks

The African green monkey kidney cells Vero (ATCC CCL-81™) and Vero E6 (ATCC CRL-1586™) were used to produce WNV and USUV stocks, respectively. Cells were maintained in Minimum Essential Medium (Gibco™, New York, USA) supplemented with 10% fetal bovine serum (FBS). Baby hamster kidney (BHK) cells (ATCC CCL-10™) were used for plaque assays, maintained in Dulbecco’s Modified Eagle Medium (DMEM) with 10% FBS.

A WNV lineage 2 strain (EMC/WNV/20TV2584/NL) was obtained from the European Virus Archive -Global (EVAg). This strain was isolated in 2020 from the common chiffchaff (*Phylloscopus collybita*) in Utrecht, the Netherlands. Vero cells were used to produce a WNV passage 4 stock for mosquito infections. Two USUV strains were used in this study: USUV/SE/17 Europe 3 lineage (USUV EU, Genbank: MK230892) and the USUV/GR/17 Africa 3 lineage (USUV AF, Genbank: MK230891) (9). Both strains were isolated in 2017 from Eurasian blackbirds (*Turdus merula*) in the province of Liège, Belgium. USUV stocks for mosquito infections were produced on Vero E6 cells after 4-6 passages.

### Oral Infection

Batches of unfed female mosquitoes were placed in paper cups and transported to a Biosafety Level 3 facility. Females were sugar-starved from 12-48 hours prior to blood-feeding and kept in an incubator maintained at 25°C and 70% RH without light, to simulate nighttime. During the evening, a Hemotek® feeding system (Hemotek, Blackburn, UK) was used to deliver an infectious bloodmeal consisting of a 2:1 mixture of chicken blood and FBS, 5 mM adenosine triphosphate (ATP), and virus stock. The final infectious titer in the bloodmeal was 1.0 x 10^7^ TCID_50_/ml WNV or USUV, representative of viremic titers in infected birds (40). Females were allowed to feed for maximum 1 hour in the dark incubator, after which mosquitoes were sedated and sorted over dry ice. Blood-fed and unfed females were separated into individual cardboard cups and provided with 10% sucrose solution *ad libitum* in an incubator maintained at 25°C and 70% RH and a photoperiod of 16:8 light:dark hours (40). Blood-fed females were held for an incubation period of 14 days, while unfed females were kept until the following oral infection. Unfed females that did not feed during a second (identical) feeding attempt were safely discarded.

### Salivation & Dissection

Mosquitoes were sugar-starved 24 hours prior to salivation. At 14 days post-infection, mosquitoes were sedated over dry ice and their wings and legs were removed using forceps. The wings and legs of each mosquito were placed in 300 µl of phosphate buffered saline (PBS) in homogenate tubes with 2.8 mm Precellys® ceramic beads (Bertin Technologies, Montigny-le-Bretonneux, France). To collect saliva, the proboscis of each mosquito was placed for 1-1.5 hours in a 20 µl pipette tip containing a 1:1 mixture of FBS and 50% sucrose solution (26). Each saliva sample was then diluted in an Eppendorf tube containing 40 µl DMEM with 5% HEPES (26). The mosquito heads were dissected using fine sterile forceps and placed in the same homogenate tubes as their respective wings and legs. The mosquito bodies were placed in new homogenate tubes containing 600 µl of PBS. All samples were stored at −80°C until further use.

### Infection Assessment

The presence of infectious virus in mosquitoes was determined by plaque assay. Mosquito bodies were homogenized using a Precellys® Evolution homogenizer at 4,500 rpm for 1 min. The homogenate was centrifuged at 13,000 rpm for 1 min (MegaStar® 1.6R, VWR International, Radnor, USA) and the supernatant was transferred to an Eppendorf tube with a 0.8 µm filter and filtered at 13,000 rpm for 3 minutes. Mosquito heads were homogenized at 6,800 rpm for 1 min. These homogenates were spun down for 1 min at 8,000 rpm and the supernatants were filtered through a 0.8 µm filter at 10,000 rpm for 2 minutes. The body samples will be further processed for viral metagenomic analysis (unpublished data by De Coninck et al.), which is why they were homogenized and spun differently from the head samples. The head or body filtrates and the saliva suspensions were added to individual wells of a 24-well plate pre-seeded with BHK cells in DMEM with 2% FBS and 1% 100 U/ml penicillin & streptomycin (PenStrep). After 2 hours of incubation at 37°C, the inoculum from the wells was removed and replaced with 0.8% carboxymethylcellulose (CMC) agar. After 3 days (for saliva samples) or 5 days (for the body and head, wing, and leg samples) of incubation at 37°C, the cells were fixed with 3.6% paraformaldehyde and dyed with crystal violet to observe plaques. Saliva samples were incubated for 3 days, allowing for smaller plaques that are more easily countable to calculate plaque forming units (PFU) per saliva sample.

The presence or absence of WNV or USUV RNA in the bodies and heads, wings and legs was confirmed using qRT-PCR. RNA extraction was performed using the NucleoSpin RNA Virus Kit (Macherey-Nagel, Düren, Germany) following the manufacturer’s instructions. WNV detection was performed by qRT-PCR amplifying the 3’UTR region using a primer pair and probe described elsewhere (41) with the probe modified to a double-quenched probe (5’-6-FAM/CTCAACCCC/ZEN™/AGGAGGACTGG-IABk®FQ-3’; Integrated DNA Technologies, Coralville, USA). For each reaction, a 20 µl mixture containing 3 µl of RNA was prepared using the Low ROX One-Step qRT-PCR 2X MasterMix kit (Eurogentec®, Seraing, Belgium) following the manufacturer’s instructions. The cycle program included reverse-transcription (48°C, 30 minutes) and incubation (95°C, 10 minutes) followed by 40 amplification cycles with denaturation (95°C, 15 seconds) and annealing (55°C, 1 minutes) steps. A qRT-PCR for detection of the USUV NS5 gene was performed using primers designed previously (42) and a modified probe sequence (5’-FAM-TGGGACACCCGGATAACCAGAG-TAMRA-3’). For each reaction, a 25 µl reaction mixture with 3 µl of RNA was prepared using the same kit and cycle conditions described above, with the exception of a 60°C annealing temperature (42). The WNV and USUV genome copies per sample were quantified using gBlocks™ (Integrated DNA Technologies Inc., Coralville, USA).

### Species Identification

DNA extraction of mosquito bodies was performed using the QIAmp DNA Mini Kit (Qiagen, Hilden, Germany) following the manufacturer’s instructions. The subspecies *Culex pipiens pipiens*, *Culex pipiens molestus*, and *Culex pipiens-molestus* hybrids were distinguished by a duplex qRT-PCR targeting the CQ11 microsatellite region. The primer pair was universal to both biotypes (43) while the probes were biotype-specific to either *Culex pipiens pipiens* (43) or *Culex pipiens molestus* (44). Hybrid *Culex pipiens-molestus* were detected by the presence of amplification curves from both probes. A 25 µl reaction volume was prepared for each reaction using the Low ROX One-Step qRT-PCR 2X MasterMix kit (Eurogentec®, Seraing, Belgium) following the manufacturer’s instructions. The cycle conditions included an initial denaturation step at 95°C for 10 minutes, 40 cycles of denaturation at 94°C for 40s, elongation at 48°C for 1 minute, and extension at 72°C for 1 minute, and a final hold stage at 72°C for 2 minutes.

Other species were identified by sequencing the cytochrome oxidase 1 (COX1) mitochondrial gene. From the DNA extractions, a 710 bp region was amplified by PCR using previously described primers (45) and the KAPA HiFi HotStart ReadyMix PCR Kit (Roche, Basel, Switzerland) following the manufacturer’s instructions. The PCR product was run on a 2% agarose gel using gel electrophoresis and the band was purified with the Wizard® SV Gel and PCR Clean-Up System (Promega, Madison, USA). Samples were submitted to Macrogen Europe (Amsterdam, the Netherlands) for Sanger sequencing. The obtained sequences were trimmed and assembled with BioEdit© v7.2.5 to produce a single consensus sequence per mosquito. Using NCBI BLASTn, the consensus sequences were compared to the standard nt database to identify the closest related hits with >99% sequence similarity.

### Wolbachia Detection

The presence of *Wolbachia* was detected using PCR on DNA extracts of mosquito bodies. The universal *Wolbachia* primers 81F and 691R amplifying the *wsp* gene were used for general detection of supergroups A and B, as described elsewhere (46). GoTaq® Green Master Mix (Promega) was used following the manufacturer’s protocol to prepare a 20 µL reaction mix containing 0.5 µM of each primer and 5 µL of template DNA. The thermocycler conditions were: initial denaturation at 95°C for 2 minutes; 35 cycles of denaturation at 95°C for 45 seconds, annealing at 50°C for 2 minutes, and extension at 72°C for 1 minute; and a final extension step at 72°C for 5 minutes. Amplified products were electrophoresed on a 2% agarose gel. A subset of PCR fragments was purified with the Wizard® SV Gel and PCR Clean-Up System and sequenced by Macrogen Europe to confirm the correct amplification target.

### Data Analysis & Presentation

A map of the collection sites was produced in QGIS v3.18.3 [QGIS Development Team (2021). QGIS Geographic Information System. Open Source Geospatial Foundation Project. http://qgis.osgeo.org]. All figures and statistical analyses were generated with GraphPad Prism v9.5.1 (GraphPad Software, San Diego, California USA). Infection rate (IR) was calculated as the proportion of blood-fed mosquitoes with infectious virus present in the body; dissemination rate (DR) was the proportion of mosquitoes with infectious virus in the body as well as in the heads, wings and legs; and transmission rate (TR) was the proportion of mosquitoes with a disseminated infection that also had infectious virus present in the saliva. Viral genome copies were statistically compared using the Mann-Whitney *U* test and the effect of *Wolbachia* infection on virus infection rate was determined using the Fisher’s exact test. A p-value of <0.05 was considered statistically significant.

## Results

### Mosquito Collections in Leuven, Belgium

A total of 1,951 *Culex* mosquitoes were captured over 166 trap nights (Fig. 2A-B). Most mosquitoes were collected at the urban site (n=13 mosquitoes/trap night) followed by the peri-urban site (n=10 mosquitoes/trap night). The majority of all captured mosquitoes were female (58.3%, n=1,137), of which 44.9% (n=876) were unfed, 12.7% (n=248) were gravid, and 0.7% (n=13) were engorged with blood.

**Figure 2.**
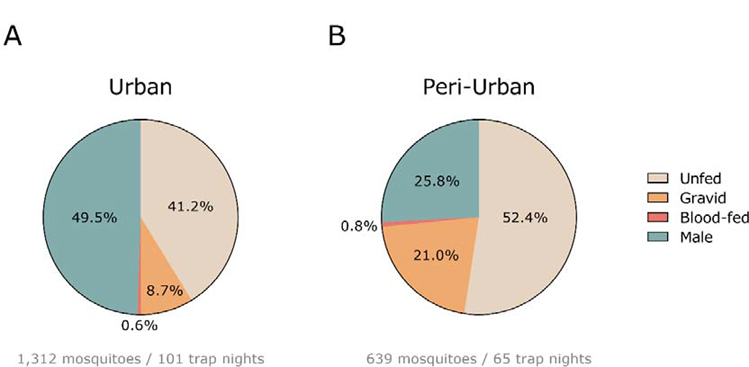
Adult *Culex* mosquitoes captured at the urban (A) and peri-urban (B) sites in Leuven, Belgium.

### Blood-Feeding & Species Identification

A total of 475 unfed females were offered an infectious bloodmeal containing either WNV (lineage 2, Netherlands 2020), USUV Europe strain (EU, lineage 3, Belgium 2016), or USUV Africa strain (AF, lineage 3, Belgium 2016). The overall blood-feeding rate was 37.9% (n=180; 95% CI: 42.3-33.6) with a 14-day post-feeding mortality rate of 14.4% (n=26; 95% CI: 20.3-10.1). Mosquito body, head, wings and legs, and saliva samples from 154 females were harvested at 14 days post-infection (Fig. 3). The majority of mosquitoes were identified as *Culex pipiens pipiens* (90.9%, n=140), dispersed evenly among the three infection groups. Eleven mosquitoes (7.1%) were identified as *Culex modestus,* belonging to the WNV (n=6) and USUV AF (n=5) infection groups. No *Culex modestus* were present in the group fed with USUV EU. The remaining mosquitoes were identified as *Culex pipiens molestus* (0.6%, n=1) in the WNV group, and a *Culex pipiens-molestus* hybrid (0.6%, n=1) and *Culex torrentium* (0.6%, n=1) in the USUV EU group.

**Figure 3.**
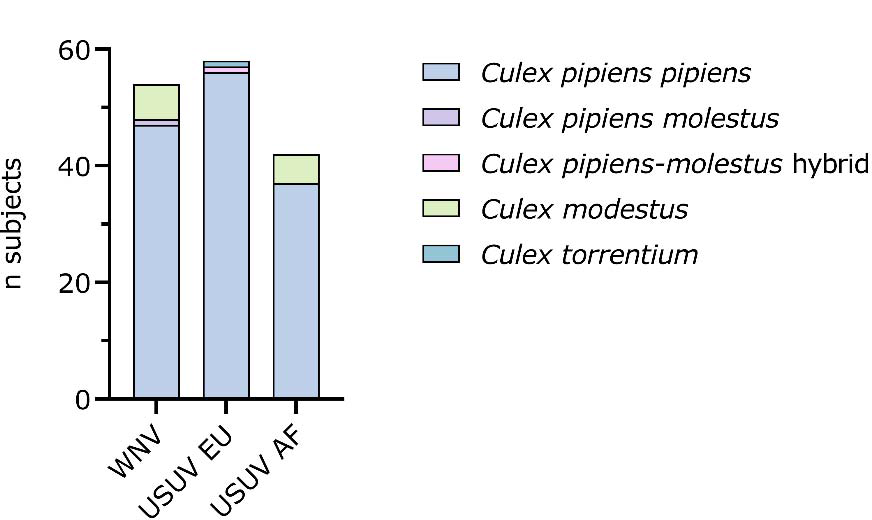
***Culex* species identification by virus infection group.** *Culex* mosquitoes were identified morphologically to the genus level, after which *Culex pipiens* biotypes (*pipiens*, *molestus*, *pipiens-molestus* hybrids) were identified by qRT-PCR while other species (*Culex torrentium* and *Culex modestus*) were identified by sequencing the cytochrome oxidase 1 (COX1) gene.

### Vector Competence

Belgian *Culex pipiens pipiens* were found to transmit WNV, but not USUV (Fig. 4A). The infection rate, as determined by plaque assay, for WNV-fed mosquitoes (n=47) was 10.6% in the bodies, followed by 40% dissemination to the head, wings, and legs and 100% transmission in the saliva. For *Culex pipiens pipiens* fed with USUV EU (n=56), 12.5% had a positive infection in the body, but there was no disseminated infection. The infection rate for *Culex pipiens pipiens* blood-fed with USUV AF (n=37) was 16.2% with a dissemination rate of 16.7%, but there was no detectable virus in the saliva.

**Figure 4.**
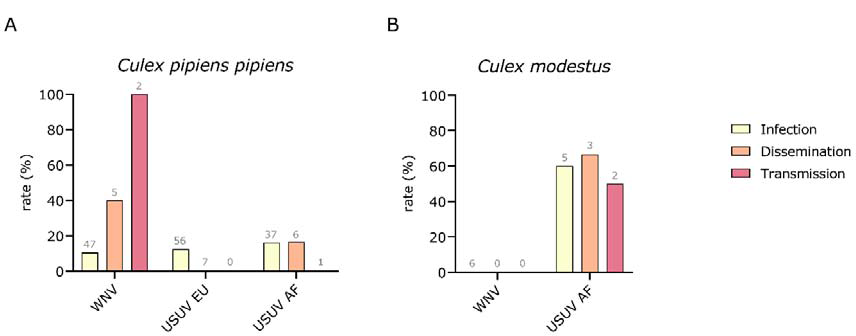
Vector competence of *Culex pipiens pipiens* (A) and *Culex modestus* (B) for WNV and USUV. The bars represent the rates of infection in the body (yellow), dissemination to the head, wings, and legs (orange), and transmission potential in the saliva (red) as determined by plaque assay. Grey labels indicate the total number of mosquitoes.

Interestingly, *Culex modestus* was the only species able to transmit USUV (Fig. 4B). A positive USUV AF infection was observed in the body in 60% of the mosquitoes tested (n=5), with 66.7% dissemination to other organs and 50.0% transmission from saliva. In contrast, there was no infection in *Culex modestus* that received a blood meal containing WNV (n=6). *Culex pipiens molestus* (n=1) was negative for WNV, and the *Culex pipiens-molestus* hybrid (n=1) and *Culex torrentium* (n=1) were both negative for USUV EU.

### Virus Quantification

The mean WNV titer in *Culex pipiens pipiens* was 1.37 x 10^8^ (95% CI: 0.0-3.35 x 10^8^) and 3.46 x 10^7^ (95% CI: 0.0-3.07 x 10^8^) genome copies per body and head, respectively (Fig. 5A-B). Females infected with USUV EU had a mean titer of 4.49 x 10^6^ (95% CI: 0.00-9.00 x 10^6^) genome copies per body, with no quantifiable RNA in the head, wings and legs. One *Culex pipiens pipiens* female with a positive plaque assay for the body but no detectable plaques for the head, wings and legs sample had quantifiable USUV AF RNA in the head, wings and legs (2.8 x 10^3^ genome copies). Including this mosquito, the *Culex pipiens pipiens* infected with USUV AF had a mean body titer of 4.78 x 10^6^ (95% CI: 4.22 x 10^4^-9.51 x 10^6^) genome copies and mean head titer of 6.06 x 10^5^ (95% CI: 0.00-8.28 x 10^6^) genome copies. There was no significant difference in viral genome copies between *Culex pipiens pipiens* bodies infected with USUV EU or USUV AF (p=0.6282), but WNV-infected bodies had significantly higher genome copies than those infected with USUV EU (p=0.0303) or USUV AF (p=0.0303). The WNV and USUV AF titers in *Culex pipiens pipiens* heads were not significantly different (p=0.3333).

**Figure 5.**
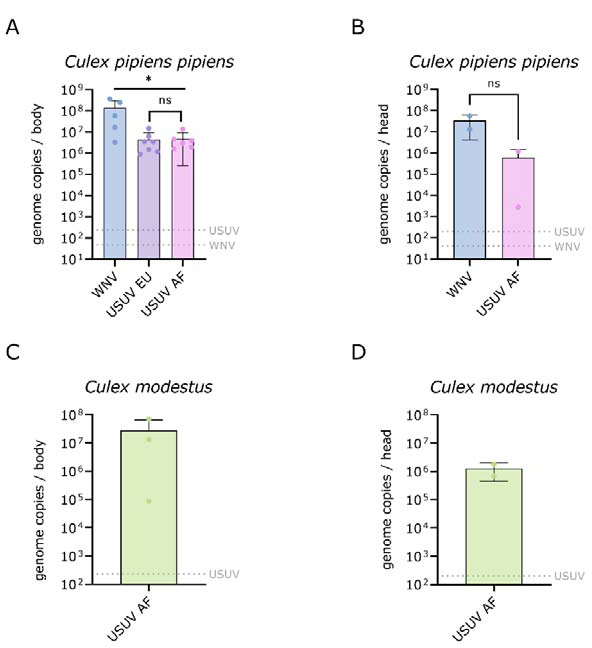
Viral genome copies in the bodies and heads of *Culex pipiens pipiens* (A-B) and *Culex modestus* (C-D). The bars show the mean viral genome copies ± SD; the grey dotted lines represent the limit of detection (LOD) of the qRT-PCR assays used.

*Culex modestus* infected with USUV AF (Fig. 5C-D) had a mean titer of 2.80 x 10^7^ (95% CI: 0.00-1.21 x 10^8^) genome copies in the body and a mean titer of 1.25 x 10^6^ (95% CI: 0.00-8.40 x 10^6^) genome copies in the head, wings and legs. There was no significant difference in USUV AF genome copies between the bodies (p=0.71) or head, wings and legs (p=>0.99) of infected *Culex pipiens pipiens* and *Culex modestus* (Fig. S1).

The mean infectious titer in *Culex pipiens pipiens* with detectable WNV in the saliva was 305±127 PFU per sample (Fig. 6). The single *Culex modestus* with a transmissible USUV AF infection had 72 PFU per sample.

**Figure 6.**
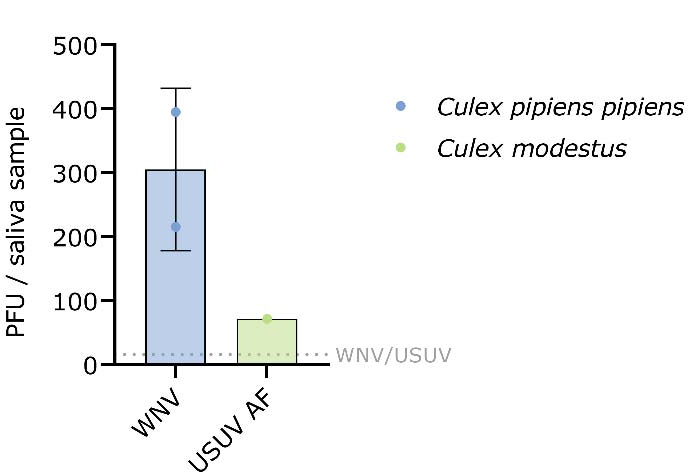
**Infectious titer per mosquito saliva sample**. Infectious virus titers were determined by plaque assay (PFU; plaque forming units); the grey dotted line represents the limit of detection (LOD) of the qRT-PCR assays used.

### *Wolbachia* Infection of Belgian *Culex* mosquitoes

All *Culex pipiens pipiens* body samples (n=139), except for one female blood-fed with WNV but with no detectable WNV infection in the body or head, wings and legs, were positive for the *Wolbachia wsp* gene (Table 1). All *Culex modestus* mosquitoes (n=11) were negative for *Wolbachia*, regardless of their infection status for USUV AF. The individual *Culex pipiens molestus* and *Culex pipiens-molestus* hybrid were both infected with *Wolbachia*, whereas the single *Culex torrentium* was not. There was no correlation between *Wolbachia* infection on WNV or USUV AF infection, dissemination, or transmission rates in *Culex pipiens pipiens* (p=>0.99) or *Culex modestus* (p=>0.99), respectively (Table S1).

**Table 1.** Prevalence of *Wolbachia* infection per species by detection of the *wsp* gene.

| Species | % *wsp* positive (n) | % *wsp* negative (n) |
| --- | --- | --- |
| *Culex pipiens pipiens* | 99.3 (139) | 0.7 (1) |
| *Culex modestus* | 0 | 100(11) |
| *Culex pipiens molestus* | 100 (1) | 0 |
| *Culex pipiens-molestus* hybrid | 100 (1) | 0 |
| *Culex torrentium* | 0 | 100 (1) |

## Discussion

We present the first report that field-collected Belgian *Culex* mosquitoes can transmit WNV and USUV in a laboratory setting. Interestingly, despite *Culex pipiens pipiens* being generally considered a USUV vector, they were unable to transmit USUV from two different strains isolated in Belgium (Europe lineage 3 and Africa lineage 3). This is in line with other studies on field-collected *Culex pipiens* mosquitoes. In a UK surveillance study, no USUV RNA could be detected in pooled samples comprising 4,800 *Culex pipiens sensu lato* (s.l.) mosquitoes (47). A field study on French *Culex pipiens* s.l. also observed a low infection rate for USUV EU lineage 3 (1.4%) while infection was much higher for WNV lineage 1 (38.7%) (48). Furthermore, a study on American *Culex pipiens pipiens* found that they were unable to transmit USUV (a Netherlands 2016 isolate) (49). Other investigations of USUV in European *Culex pipiens pipiens* were performed with laboratory-colonized mosquitoes (24,25) or did not specify the *pipiens* and *molestus* biotypes (7,15,57–66,27,50–56). It is therefore possible that the true USUV vector competence of *Culex pipiens pipiens* is significantly lower in nature than what is measured in lab colonies. The midgut escape barrier and salivary gland infection barrier may be key in preventing USUV transmission in *Culex pipiens pipiens*, since USUV EU and AF were able to establish in the midgut and/or disseminate to the rest of the mosquito but not reach the salivary glands. On the other hand, Belgian *Culex pipiens pipiens* proved to be efficient vectors of WNV.

In contrast to the lack of USUV transmission by *Culex pipiens pipiens*, the *Culex modestus* captured in this study were competent vectors for USUV AF. This result was unexpected, given that the sample size of *Culex modestus* in the USUV AF group was low (n=5). The small sample size, especially compared to the number of *Culex pipiens pipiens* tested in this study (n=140), suggests that *Culex modestus* is a highly efficient USUV AF vector. USUV RNA has previously been detected in pooled *Culex modestus* from the Czech Republic (14) but, to our knowledge, our study is the first to demonstrate USUV vector competence in *Culex modestus* using live mosquitoes. Conversely, no *Culex modestus* in the WNV-fed group (n=6) developed a WNV infection, which contrasts with vector competence studies on laboratory colonies that suggest that *Culex modestus* is an efficient vector for WNV (12,28). There is also evidence of WNV RNA detected in field-captured *Culex modestus* from Italy (27). Therefore, it is possible that our sample size was not high enough to obtain WNV-infected *Culex modestus*.

The viral RNA copies quantified in the mosquitoes in this study are consistent with other reports that measured viral loads in *Culex pipiens pipiens* infected with WNV (48,67) and USUV (67). There was no significant difference in *Culex pipiens pipiens* body titers between USUV EU and USUV AF, but only USUV AF was able to disseminate to the head, wings, and legs. Similarly, there was no significant difference between USUV AF genome copies between *Culex pipiens pipiens* and *Culex modestus* bodies or heads, yet *Culex modestus* was the only species with detectable USUV AF in the saliva. It has been demonstrated elsewhere that RNA copies do not necessarily correlate with the quantity of infectious virus or the ability to establish persistent infection or dissemination in the mosquito (68). Our results suggest that the USUV AF lineage 3 replicates more efficiently in *Culex modestus* than in *Culex pipiens pipiens*. Furthermore, USUV AF may be more efficient than USUV EU in bypassing the midgut barrier and/or host immune response of *Culex pipiens pipiens*. These findings are especially interesting, as most cases of USUV isolated from avian samples in Belgium belonged to the EU lineage (9). Furthermore, in a study using the same USUV strains as this study, the EU strain produced higher quantities of viral RNA than the AF strain when inoculated in chicken embryo-derived cells (69). More research is thus needed to understand the vector competence of *Culex modestus* to USUV EU strains.

As research interest in *Wolbachia* continues to grow due to its success as an arbovirus control strategy, we determined the prevalence of *Wolbachia* infection in the mosquitoes challenged in this study. Almost all *Culex pipiens pipiens* had a *Wolbachia* infection (99.3%), consistent with other European studies (32,37). The only *Culex pipiens pipiens* without *Wolbachia* was in the WNV-fed group with no detectable WNV infection. We did not find a statistically significant effect of *Wolbachia* infection on the ability of WNV or USUV to replicate in *Culex pipiens pipiens*; however, we emphasize that there were not sufficient Wolbachia-negative mosquitoes to reach an accurate conclusion. A limitation to our study is that we did not quantify *Wolbachia* loads; however, a study on *Culex pipiens pipiens* from Germany found no correlation between *Wolbachia* levels and WNV infection (32). Of the *Culex modestus* identified in this study, all were negative for *Wolbachia*. In contrast to our findings, other studies have found *Wolbachia* in *Culex modestus* from Italy (prevalence rate unknown) (38) and Eastern Europe (7% prevalence in pooled samples) (39). However, our sample size was likely too low to reach the conclusion that Belgian *Culex modestus* do not carry *Wolbachia*. As almost all *Culex pipiens pipiens* were positive for *Wolbachia* but all *Culex modestus* were negative, we can presume that the probability of acquiring and maintaining *Wolbachia* is species-dependent. It would be interesting to investigate if the presence of *Wolbachia* in *Culex modestus*, whether acquired naturally or artificially, plays a role in their vector competence.

In this study, field-captured mosquitoes were the preferred model of choice to study vector competence over laboratory colonies. Multiple intrinsic and extrinsic factors can influence the fitness and vector competence of mosquitoes, such as genetic diversity, age, parity rate, the microbiome, innate immunity, climate, and the environment (70). It has also been shown that vector competence can differ significantly between wild and lab-colonized mosquitoes of the same species (20,71). Despite the advantages of using field mosquitoes over colonies, there are several limitations, such as the dependence on climate, species identification, mosquito loss or damage during collections, and difficulty in achieving high mosquito numbers. A limitation of our study was that all *Culex* species were pooled for oral infections prior to species identification, which is why we did not obtain any *Culex modestus* fed with USUV EU. However, despite the evident drawbacks, we argue that field mosquitoes provide a more accurate measure of vector competence because they represent the natural vector population.

To date, blood and organ donor screenings for WNV take place in endemic European countries, but for the prevention of WNV and USUV transmission by mosquitoes there is little routine vector control in place. In areas where competent vectors are present, it is important to determine the risk of transmission by measures of vector competence and vectorial capacity to anticipate and prepare for potential outbreaks. We present the first evidence that *Culex* mosquitoes from Belgium can transmit WNV and USUV. *Culex pipiens pipiens* were capable of transmitting WNV, whereas *Culex modestus* was shown to be an efficient vector for USUV. More research is needed to understand if *Culex modestus* can transmit other USUV lineages, such as EU lineage 3, and if *Wolbachia* can influence the vector competence of *Culex modestus*. As Belgium lies between countries with prior human WNV cases and is known to be endemic for USUV, and add to that the presence of competent mosquito vectors for both arboviruses, it is not unlikely that Belgium will experience human cases of WNV and USUV in the future. Improved mosquito surveillance and arbovirus prevention measures in Belgium are therefore highly recommended.

## Acknowledgments

We would like to thank Prof. Mutien Garigliany from the University of Liège for providing us with the Belgian USUV stocks. We would like to recognize Jan Vandyck and the kind workers at the Botanical Garden (Kruidtuin) of Leuven as well as Marc Leroy and members of the Space and Real Estate Division at KU Leuven (Arenberg Park) for kindly allowing us to set mosquito traps. We would like to thank Dr. Suzanne Kaptein and Elke Maas for sharing their WNV qRT-PCR protocol with us, and we thank Joske Depovere for her assistance with the mosquito collections. Finally, we thank Prof. Johan Neyts for allowing us to use his laboratory space and equipment.

## Notes

### Competing Interest Statement

The authors have declared no competing interest.

