## Supplemental material for "Belgian *Culex pipiens pipiens* are competent vectors for West Nile virus but not Usutu virus"

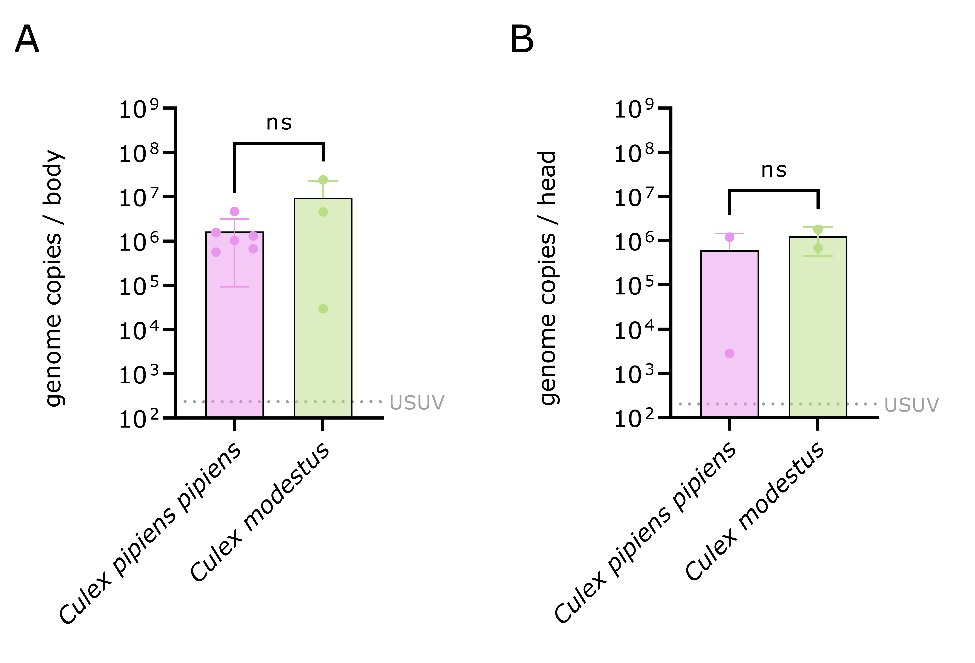

### Figure S1. Comparison of USUV AF genome copies in *Culex pipiens pipiens* and *Culex modestus*. USUV AF genome copies were determined by qRT-PCR in individual bodies (A) and heads (B). The bars show the mean viral genome copies ± SD; the grey dotted lines represent the limit of detection (LOD) of the qRT-PCR assays used. Statistical analysis was performed with the Mann-Whitney test.

### Table S1. Impact of *Wolbachia* infection on virus infection, dissemination, and transmission by species. The effect of *Wolbachia* infection on arbovirus infection rate (IR), dissemination rate (DR), and transmission rate (TR) were determined by the Fisher’s exact test. NS: non-significant.

| **Species** | **Effect on IR, p-value** |  | **Effect on DR, p-value** | **Effect on TR, p-value** |
| --- | --- | --- | --- | --- |
| *Culex pipiens pipiens* | NS, p=>0.9999 |  | NS, p=>0.9999 | NS, p=>0.9999 |
| *Culex modestus* | NS, p=>0.9999 |  | NS, p=>0.9999 | NS, p=>0.9999 |
